# CLADES: A Classification-based Machine Learning Method for Species Delimitation from Population Genetic Data

**DOI:** 10.1101/282608

**Authors:** Jingwen Pei, Chong Chu, Xin Li, Bin Lu, Yufeng Wu

## Abstract

Species are considered to be the basic unit of ecological and evolutionary studies. Since multi-locus genomic data are becoming increasingly available, there has been considerable interests in the use of DNA sequence data to delimit species. In this paper, we show that machine learning can be used for species delimitation. There exists no species delimitation methods that are based on machine learning. Our method treats the species delimitation problem as a classification problem. It is a problem of identifying the category of a new observation on the basis of training data. Extensive simulation is first conducted over a broad range of evolutionary parameters for training purpose. Each pair of known populations are combined to form training samples with a label of “same species” or “different species”. We use Support Vector Machine (SVM) to train a classifier using a set of summary statistics computed from training samples as features. The trained classifier can classify a test sample to two outcomes: “same species” or “different species”. Given multi-locus genomic data of multiple related organisms or populations, our method (called CLADES) performs species delimitation by first classifying pairs of populations. CLADES then delimits species by maximizing the likelihood of species assignment for multiple populations. CLADES is evaluated through extensive simulation and also tested on real genetic data. We show that CLADES is both accurate and efficient for species delimitation when compared with existing methods. CLADES can be useful especially when existing methods have difficulty in delimitation, e.g. with short species divergence time and gene flow.

## 1 Introduction

Species are considered to have fundamental importance in ecological and evolutionary studies. In the light of new sequencing technologies, the use of multi-locus sequence data and identification of species status have undergone a revolution during the last two decades (Rannala, 2015). Many papers have been written on, and various computational approaches have been developed for, species delimitation (e.g. Yang and Rannala, 2010; Ence and Carstens, 2011; Jones et al., 2014). However, there has been significant confusion on species delimitation (Rannala, 2015; Carstens et al., 2013). One of the main issues is the lack of the widely acceptable definition of the concept of species itself. It is reported that there have been over 20 definitions of species (Wilkins, 2009; Hausdorf, 2011). For example, Biological Species Concept (BSC) defines a species as populations that actually or potentially interbreed in nature (Mayr, 1976). Another example is the evolutionary species concept (ESC), which defines a species to be a lineage that maintains its identity from other lineages and has its own evolutionary tendencies and historical fate (Wiley, 1978). Bayesian approaches such as BP&P (Yang and Rannala, 2010) often rely on ESC. Clearly, it is difficult to quantify one concept to a well-defined “delimiter” (Rannala, 2015). The difficulty in specifying the species concept makes it even harder to develop computational methods for species delimitation.

Despite the difficulty in defining species, there has been active research for developing computational approaches for species delimitation (e.g. Rannala and Yang, 2013; Ence and Carstens, 2011; Jones et al., 2014; Birky Jr, 2013). Recently, model-based methods (e.g. Bayesian approaches) for species delimitation with multi-locus sequence data became popular in the literature. The Bayesian approach is usually based on the multi-species coalescent (MSC) model (e.g. Takahata et al., 1995; Rosenberg et al., 2002; Rannala and Yang, 2003). The representative approach for Bayesian species delimitation is the BP&P approach (Yang and Rannala, 2010; Rannala and Yang, 2013). Other recently developed methods for species delimitation include BFD* (Leache et al., 2014) and PHRAPL (Jackson et al., 2017).

In this paper, we take the evolutionary species concept (ESC). That is, populations are called species if they are isolated by evolutionary timescale. Here, evolutionary timescale mainly refers to two quantities:

1. Divergence time *τ* of two species or populations.
2. The per generation per individual migration rate *m* between two species or populations.

Now imagine we consider two groups of gene sequences that are sampled from two populations. As biologists, we want to know whether: 1) these distinct genetic groups are actually two species that diverged *τ* coalescent units ago and exchange migrants at rate *m* after the population split, or these genetically distinct populations belong to the same species. That is, the problem of species delimitation we consider in this paper is essentially a statistical inference problem. Assume for now the mutation rate per generation per base *µ* is fixed. Suppose the two groups are indeed from two distinct species. Intuitively, when *θ* = 4*Nµ* (*N* is effective population size) and *τ* decreases (i.e. shallow divergence times), it becomes statistically more difficult to distinguish between the two cases (Rannala, 2015; Zhang et al., 2011). When the scaled migration parameter *M* = *Nm ≤* 0.1, it virtually has no effect in determining whether the two groups are from two different species (Zhang et al., 2011). As *M* increases, the two species exchange gene sequences more often and thus it becomes more difficult to tell apart the two cases. It is stated in Zhang et al. (2011) that *M* = 1 is the isolation threshold for Bayesian methods. With high migration *M ≥* 10 per generation, species delimitation methods strongly favor one-species model. Many existing methods (e.g. BP&P) aim to classify the given gene sequences that fall within the cases specified by the evolutionary thresholds above.

While approaches such as BP&P are certainly very useful and have been used extensively in the literature, existing methods such as BP&P work less well for “harder” cases when the genetic isolation timescale is near the evolutionary threshold and when migration event involves in the evolutionary process (Sukumaran and Knowles, 2017; Rannala, 2015). Another issue is gene flow. Most existing species delimitation approaches (including BP&P and BFD*) don’t consider gene flow in their models. The impact of gene flow for species delimitation has started to be studied recently (Jackson et al., 2017). Moreover, many existing species delimitation methods (e.g. BP&P and PHRAPL) are often very slow for large data. This raises two questions: (i) Can one design more accurate and efficient methods that give more accurate species delimitation results than existing methods (e.g. BP&P) at these harder cases? (ii) Can such method also reliably find the underlying genetic isolation with gene flow?

We note that the underlying evolutionary thresholds are specific to the inference method, not a fundamental aspect of the species delimitation problem itself. That is, it is conceivable that a different computational method can statistically distinguish cases that violate the evolutionary thresholds where approaches such as BP&P cannot distinguish. Such a method can be useful, for example, if one is interested in studying very closely-related species. Rannala (2015) proposed a resolution of focusing on characteristic patterns of genetic divergence between populations for species delimitation. In this paper, we follow this general approach and explore the feasibility of detecting genetic divergence in populations to delimit species. Since the concept of species is critical to any method for species delimitation, we elaborate on the concept of species. When only sequences are given and no prior knowledge about the organisms are known, one may view species delimitation as a statistical inference problem. Suppose there are two species delimitation methods: “aggressive” and “conservative”. The aggressive method tends to call “different species” more often than the “conservative” method. By doing so, the aggressive method may have less false negatives but more false positives than the conservative method. Therefore, one may view the species boundary corresponds to the statistical error threshold that the user can accept. A machine learning approach is designed to achieve good performance of species delimitation by balancing the false positive and false negative errors.

In this paper, we develop a classification based method, called CLADES (which stands for CLAs-sification based DElimitation of Species), for species delimitation. Here are the main contributions of our method.

1. CLADES is based on a classification model trained and tested with the multi-locus sequence data. The main advantage of CLADES is that it can more accurately distinguish the “same species” and “different species” cases than Bayesian methods, especially when the genetic isolation is not very ancient. Different from many existing methods, CLADES can still give reasonable delimitation results when there is gene flow.
2. CLADES is designed to determine whether two or more groups of gene sequences belong to the same or different species. Different from many existing methods, we don’t need the user to provide guide trees (also see Jones et al., 2014) or provide priors of root age and *θ*. Our method can be applied to datasets where there are more than two populations.
3. We demonstrate that CLADES is robust against various modeling assumptions, such as various population demographic events, and different evolutionary parameters.
4. CLADES is very efficient and can scale to large and long sequence dataset. It is also flexible and can also be easily extended to accommodate new requirements.

We mainly use simulated data to validate the performance of CLADES. We also apply CLADES on real biological data to demonstrate CLADES can indeed work on real data.

## 2 Method

### 2.1 Overview

Population trees model the evolutionary history of multiple populations, where these populations may originate from different species. In this paper, populations are labeled with alphabet letters and numbers. Unless otherwise stated, populations labeled with same alphabet prefix belong to the same species. Figure 1 illustrates an example of a population tree and its corresponding species tree. The population tree shows that *A*_1_ and *A*_2_ are the two populations of the same species *A*.

**Figure 1:**
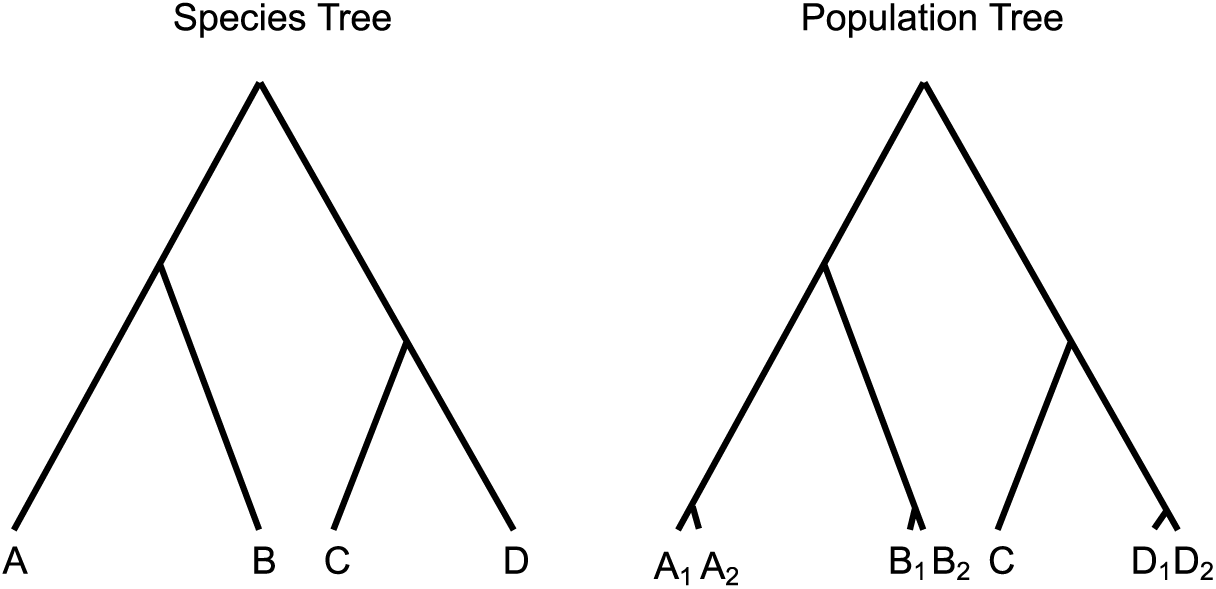
A population tree for seven populations and corresponding species tree for four species. Species A, B and D have two populations.

For a population, we collect a number of genomic sequences at multiple loci. We assume they evolve according to the standard population genetic models. In particular, mutation rate *µ* and effective population size *N* are assumed to be constant among all loci. Given multi-locus sequence data from several populations, our objective is to assign these populations to a number of species. For simplicity, we first focus on two populations. If given multiple populations, we consider each pair of populations separately and then conduct species delimitation by maximizing the likelihood of species status assignment of these multiple populations. This is described in Section 2.4.

The key idea of our method is viewing species delimitation as a classification problem. It is a problem of determining the category of an unlabeled data sample, on the basis of training data containing samples whose categories are known. Suppose there are *n*_*A*_ and *n*_*B*_ genomic sequences collected from two populations *A* and *B* at a locus. Then a label “+1” or “-1” is either known or expected to be assigned for this group of sequences, meaning population *A* and *B* belong to the “different species” or “same species”. Here, a group of sequences at one locus from two populations is considered as one data sample. With the obtained sequences from multiple loci, we are able to collect a number of data samples for two populations. These samples can be naturally classified into two clusters based on their labels “+1” and “-1”. We use standard population genetic simulation to generate training data with known labels to train a classifier, which can be later used to determine if two populations belong to the same species or not. Note that it is possible the predicted labels at different loci for two populations are not always the same. In this case, one can take majority vote of all data samples for two populations. That is based on the percentage of loci supporting a species status between two population. We further discuss this situation in Section 2.4.

### 2.2 Summary statistics as features

Using the full information of genomic sequences is difficult for model construction and estimation. Thus, we use a list of summary statistics to represent the key information about the evolutionary history on the sequences that is contained in the sequence data. That is, summary statistics are the features we use in the classification to represent data samples. The chosen features should capture aspects of the underlying genealogical history of the sequences and help to distinguish between two clusters (“same species” and “different species”). Our method CLADES uses five informative summary statistics: the proportion of private positions, folded-SFS, pairwise difference ratio, F-statistics and longest shared tract. Each of them presents a high correlation with cluster category. For completeness, in the following, we provide more details on how we compute these summary statistics. See standard population genetics books (e.g. Hartl et al., 1997; Gillespie, 2010) for more details.

#### 2.2.1 Private position

We compute the proportion of private positions among the total number of polymorphic positions found in the group of sequences. We call a position fixed in a population if all sequences have the same allele type within the population at the position. A site is called variant if it is not fixed. For *n*_*A*_ + *n*_*B*_ sequences from population *A* and *B*, we say a position is private if the position is fixed in one population but not fixed with same allele type in the other population. The more divergent two lineages are, the more private positions in the genome we expect there to be. For example in Figure 2 the position 4 is a private position because it is fixed with allele ‘A’ in the population *A* but not fixed in the population *B*. And the position 5 is also a private position since it is fixed with allele ‘A’ in population *A* and is fixed with a different allele ‘T’ in population *B*. The proportion of private positions among these sequences is 4*/*6 = 66.67% (Figure 2).

**Figure 2:**
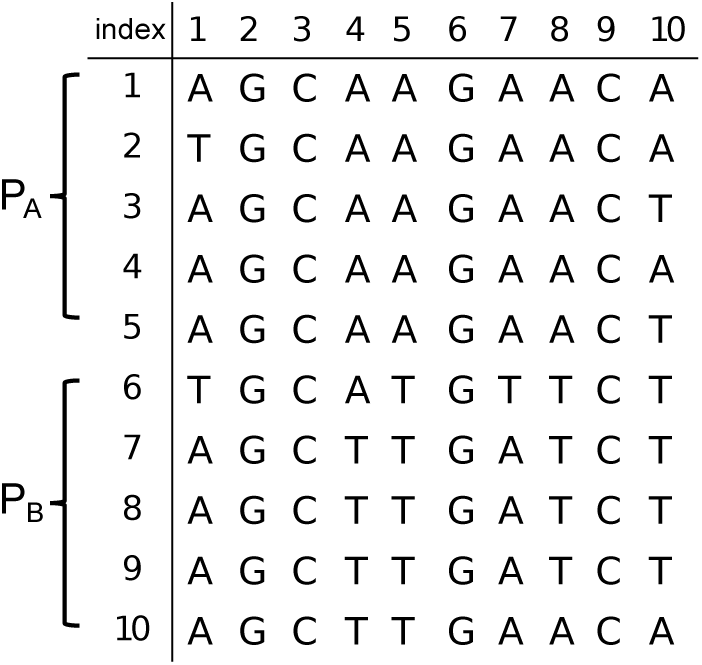
Sequences from two populations. P_A_ and P_B_ represent population A and B. Row: sequence. Column: site index.

#### 2.2.2 Folded-SFS with *k* bins

SFS (site frequency spectrum) is a vector that describes the distribution of allele frequencies over a set of sequences. Site frequency represents a different spectrum between genetic data from a single species and two different species. The *i*th element in the SFS describes the number of alleles that has *i* copies of one allele type among all *n*_*A*_ + *n*_*B*_ individuals. Here we do not consider the fixed sites. So the length of SFS is *n*_*A*_ + *n*_*B*_ − 1. The summation of SFS equals to the total number of variants *n*_*v*_. For example in Figure 2 SFS is [1, 1, 0, 2, 1, 1, 0, 0, 0]. The first element ‘1’ indicates that there is 1 allele (site 7) that has allele type ‘T’ with frequency of 1. The fourth element ‘2’ indicates there are 2 alleles (sites 4 and 8) that have allele type ‘T’ with frequency 4 among all the sequences.

Since we regard two allele types interchangeably, we further use the folded-SFS by adding the *i*th element and (*n*_*A*_ +*n*_*B*_ −*i*)-th element of SFS together to form a new vector of length (*n*_*A*_ +*n*_*B*_ −1)*/*2. For example, the folded-SFS for sequence data in Figure 2 is [1, 1, 0, 3, 1]. The 4*th* element is created by summing up 4*th* and 6*th* elements in SFS. When having a large number of individuals, SFS can be long and sparse. To make it more informative, we collapse a folded-SFS into *k* bins by dividing a folded-SFS into *k* parts from left to right with equal length and summing the elements in each part together. When having a large *k*, it might easily lose power in classification. To balance the different number of sites in different dataset, we use a unified *k* = 3 to denote three levels of spectrum as low frequency, medium frequency and high frequency sites.

#### 2.2.3 Pairwise difference ratio

Pairwise difference is useful since it measures the similarity between two populations. An existing method proposed by Birky Jr (2013) uses *K/θ ≥* 4 with 95% confidence to delimit species. Here *K/θ* is the same as the pairwise difference ratio defined here. We compute pairwise difference as the number of different alleles between two sequences. For example, sequence *AGCAAGAACA* and sequence *TGCATGTTCA* have the pairwise difference of 4 since they have the different alleles at positions 1, 5, 7 and 8. For a number of sequences, pairwise differences can be computed for pairs of sequences and the average of them can be used as summary statistics. Regarding *n*_*A*_ + *n*_*B*_ sequences from two populations, there are two types of pairwise difference means to compute: pairwise difference mean within a single population and pairwise difference mean between populations. Let *d*_*between*_ and *d*_*within*_ denote these two values respectively. We use the average of the pairwise difference mean in two populations 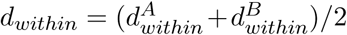 to represent the mean within population for whole sequences. For pairwise difference between populations, we compute pairwise difference mean between sequences in the population *A* and the population *B*. Pairwise difference ratio is defined as *r* = *d*_*between*_*/d*_*within*_. When two populations are distantly related, *r* would be much greater than 1.

#### 2.2.4 F-statistics *F*_*st*_

In population genetics, F-statistics, also known as fixation index, describes the level of heterozygosity of population. Suppose 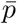 is the average frequency of an allele in all the populations combined, and 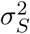 is the variance of allele frequency of each population. F-statistics is defined as 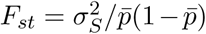 However, the quantities used in the definition are hard to estimate. In practice, we use one simple estimator of *F*_*st*_ in the following way *F*_*st*_ = (*d*_*between*_ − *d*_*within*_)*/d*_*between*_. Here *d*_*between*_ and *d*_*within*_ are defined above.

#### 2.2.5 Longest shared tract

If we view each sequence as a string, then the shared tract indicates the common substring that two sequences share. Note that the sequences are aligned, so the shared tract must be at the same position. For example in sequences 1 and 6, the region from position 2 to 4 is a shared tract and the position 6 is another shared tract. We are interested in the longest shared tract between two populations. For example, the longest shared tract length between population *A* and *B* in Figure 2 is 5 since sequences 1 and 10 are identical from position 6 to 10. Here we use the percentage of shared tract length as our summary statistics. In this case, it is 50%. The purpose of this summary statistics is to detect the migration between populations. When two populations have a high level of heterozygosity but there are long shared tracts, it is likely that these two population diverge anciently but exchange gene flow.

### 2.3 Unphased genotypes

Sometimes the given data are unphased genotypes. In this case, genotypes are known for SNP sites but the phasing information is absent. When SNP site positions are provided, it is possible to compute summary statistics from unphased genotype data. Among five types of summary statistics, folded-SFS, pairwise difference ratio, *F*_*st*_ and number of private positions can be computed accurately for both haplotype data and genotype data. The only difference occurs in the computation of longest shared tract when the genotypes have no known phased haplotypes. Here we proposed a heuristic method to compute the longest shared tract for genotypes. See Section C of the Supplementary Material for details.

### 2.4 Classification-based species delimitation

To train a classifier for delimiting species, we use a classic supervised learning approach called Support Vector Machine (SVM) (Cortes and Vapnik, 1995). Supervised learning is one type of machine learning that infers functions from training data. Here we conduct extensive simulation to collect two populations data from “different species” and “same species” under various evolutionary settings (see Section 2.7). The aim is to divide all training samples into two clusters with the least miss-classification score. As described above, each training sample can be represented as a list of summary statistics. SVM builds a support vector regression using these summary statistics and then trains the weights for each statistic iteratively to minimize the miss-classification loss. As SVM assumes the training data is in a standard range, all summary statistics are normalized to the range of [0, 1]. We use LibSVM (Chang and Lin, 2011) to conduct model training. SVM classifier can compute the probability for which cluster a training sample falls into. Suppose multi-locus sequence data are collected for two populations. The predicted labels from classifier may disagree for these two populations at different loci. Then we compute the mean of probabilities that they belong to the “same species” or “different species” across multiple loci. The species status with higher probability would be the inference result.

With the trained classifier for pairwise populations, we now build a framework based on this classifier to perform species delimitation for multiple populations. Suppose there are *n* (*n ≥* 2) populations *{P*_1_, *P*_2_, *…, P*_*n*_*}*, we first pair up every two populations and use the trained classifier to determine the species status and the associated probability for these two populations. Every pair of populations has an average probability to be assigned to “different species” or “same species” after this step. Let 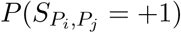 and 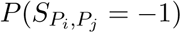 denote the probabilities that populations *i* and *j* belong to the different and same species. Suppose 𝒜 is the assignment of species delimitation for *n* populations. An assignment 𝒜 is a number of sets that contains one or multiple populations. Populations that fall into the same set belong to the same species and populations in different sets belong to the different species. For each assignment, we can compute a likelihood by multiplying the probabilities for pairwise populations 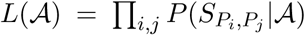 We then find the optimal assignment 𝒜* that gives maximum likelihood for the species assignments for multiple populations.

### 2.5 SNP data

Besides sequence data, CLADES can also be applied for SNP data. Given the positions for the sites, it is possible to use SNP data for species delimitation in CLADES since summary statistics can be computed as usual.

### 2.6 Cryptic sympatric species

So far we assume the number of populations and which population that samples come from are known. In practice, sometimes samples collected from the field are mixed-up. That is, samples from different species may be placed in the same group. In this case, one may want to find the cryptic sympatric species from such mixed-up samples. By clustering the samples into a number of populations, CLADES can be used for finding cryptic sympatric species (see Section D of the Supplemental Materials for details). It may be useful to consider running the clustering procedure before the CLADES analyses especially for sympatric species or even species that are believed to be allopatric. We note that many cryptic species complexes may not occupy discrete allopatric boundaries, but are instead co-distributed (i.e. sympatric). Thus, before using CLADES, grouping DNA sequences into genetically defined clusters (or populations) for all focal samples can be useful.

### 2.7 Training data generation

To obtain training data, we conduct simulation based on the two-species model (Figure 3). For simplicity, two-species model contains two species *A* and *B* that diverge at ancient time *τ* with the same population size parameters *θ*_*A*_ = *θ*_*B*_ = *θ*. Each species have two populations that split in a very recent time *τ*_*p*_. Figure 3 illustrates a population tree for the two-species model. Migration is allowed to occur between species *A* and *B* with *M* = *Nm* migrants per generation, where *m* is the migration rate per generation. With this two-species model, we use the *MCcoal* simulator (Rannala and Yang, 2003) to simulate multi-locus sequence data with length *L* under various parameters (*θ, τ, M*) for training purpose.

**Figure 3:**
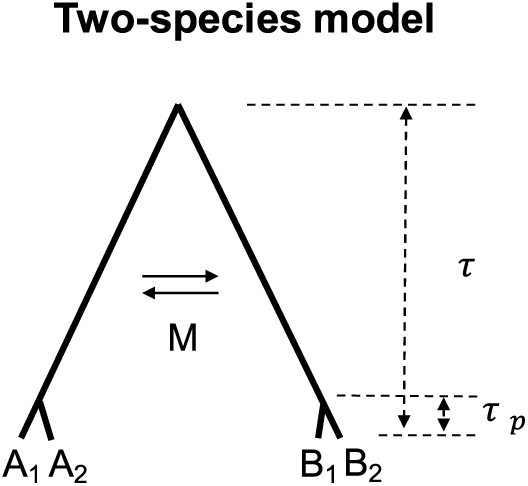
Two-species model. There are two species A and B in population tree, and each species have two populations A_1_ and A_2_, B_1_ and B_2_.

According to Zhang and Hewitt (2003), animal and plant species have a broad range of *θ ∼* (0.0005, 0.02). Thus, we choose *θ* from set *{*0.0005, 0.001, 0.002, 0.004, 0.008, 0.01, 0.02*}*. For each *θ*, we simulate two cases of species divergence time *τ* = *θ* and *τ* = *θ/*10, representing an ancient divergence and a more recent divergence between species *A* and *B*. For the case of two populations from the same species, the splitting time is fixed to *τ*_*p*_ = *θ/*500 in the model. Migration parameter *M* is chosen from the candidate set *{*0, 0.01, 0.1, 1, 3, 5*}*. For each possible (*θ, τ, M*) setting, we simulate sequences at 100 loci in *L* = 100*Kbp* for populations *A*_1_, *A*_2_, *B*_1_ and *B*_2_. And for each locus, 40 sequences are sampled, 10 for each population. For the ease of simulation, we assume symmetric migration between species *A* and *B* before populations in species split at time *τ*_*p*_.

The procedure of training a model is conducted in two steps. We first train a classifier for data simulated under each (*θ, τ, M*) setting with 4-fold cross-validation to examine the accuracy. As the training accuracy is higher than 75%, we then combine all the training samples together to train a global classifier in order to fit all possible *θ* and *M*. This way, the trained classifier doesn’t assume a fixed *θ* and *M* value. This is useful for data where these parameters are unknown. At last, we generate a small amount of test data to test the model accuracy. The amount of samples is critical in the training and testing. Large number of training samples is required to get a robust classifier.

## 3 Results

### 3.1 Accuracy of the classifier

The accuracy of CLADES mainly relies on accurate classification of data samples from two populations. Here we define the accuracy of the classifier as the percentage of accurately classified data samples (i.e. two populations data at one locus). Note that the accuracy of the classifier is different from the accuracy of the CLADES model. To evaluate the performance of the classifier, we simulate test data with various parameters (*θ, τ, M*). The candidate values of these parameters are the same as those described in Section 2.7. If not specified otherwise, for each setting we simulate sequence data at 30 loci with length *L* = 10*Kbp* in the default setting based on the two-species model.

#### 3.1.1 The impact of population size parameter *θ*

To test how the population size parameter *θ* affects the performance of the trained classifier, we combine the test data samples simulated with the same *θ* values together as a group. As shown in Figure 4 (A), the accuracy of the classifier increases as the population size parameter increases. The same is for F-score. The percentage of accurately classified samples is above 75% when *θ* is as small as 0.0005. The increase of F-score along with accuracy indicates false positives and false negatives reduce accordingly and become more balanced.

**Figure 4:**
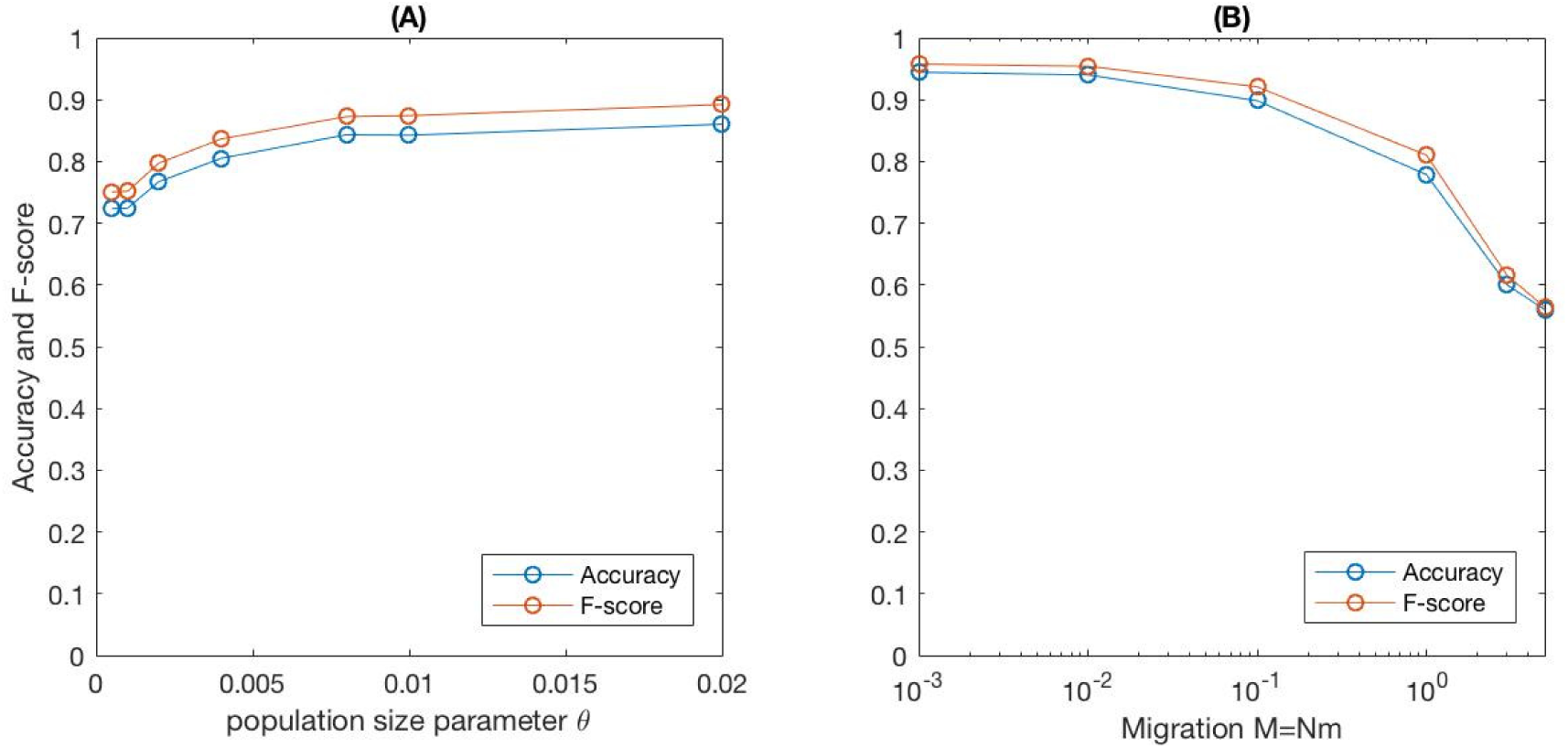
(A)The impact of population size parameter θ. (B) The impact of migration parameter M. For better visualization, x-axis is shown in logarithmic scale. Note that M = 0 cannot be plotted in logarithmic scale, thus we use 0.001 to represent M = 0 here.

#### 3.1.2 The impact of migration parameter *M*

We study the impact of the migration parameter *M* by evaluating the accuracy for each candidate *M*. Here we combine the data samples simulated with the same *M* as a group. Figure 4 (B) shows that the percentage of accurately classified samples decreases as the migration parameter increases. Recall the isolation threshold *M* = 1 is proposed by Zhang and Hewitt (2003) for Bayesian methods. We also observe a significant decrease in accuracy when *M* is greater than 1. However, the classifier can still obtain a reasonably high accuracy of 83% when *M* = 1. As *M* increases to 3 or even 5, the accuracy of *∼* 60% indicates that CLADES is still capable to distinguish two species using multi-locus data even when *M* is relatively large.

#### 3.1.3 Robustness of classifier

To further evaluate the robustness of the trained classifier, test data are simulated under different scenarios other than varying *θ* and *M*, such as one-way migration and bottleneck in population history.

The accuracy of the classifier largely relies on the stability of summary statistics. Intuitively, the variance of the computed summary statistics for two populations becomes larger when using sequence data with shorter length. This could lead to less accurate classification. Here we fix parameters *θ* = *τ* = 0.001 and *M* = 1, and then simulate multi-locus data with different lengths from *L* = 2*Kbp* to *L* = 100*Kbp*. Figure 5 (A) shows that the classification accuracy first increases linearly and then asymptotes as the simulated sequence length increases. It indicates that less samples are misclassified when using longer sequences for delimitation. Thus, in general longer sequences are preferable for species delimitation.

**Figure 5:**
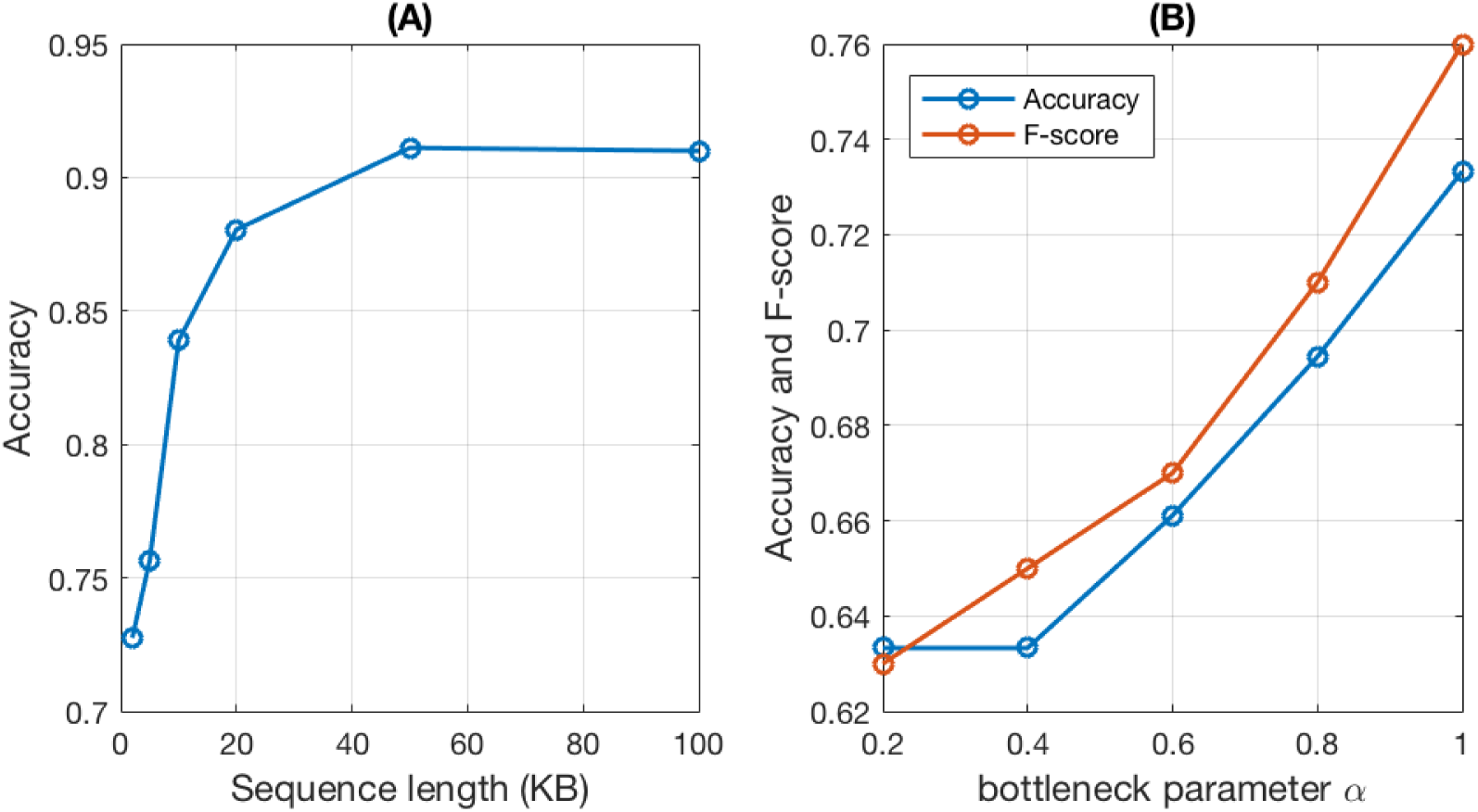
(A) Impact of sequence length, L is chosen from set {2, 5, 10, 20, 50, 100} Kbp. (B) Impact of bottleneck parameter α, N = αN_o_. N_o_ is the effective population size before and after bottleneck. Simulation parameters: θ = 0.01, τ = 0.01, M = 1.

In the two-species model, migrations between two species *A* and *B* are assumed to occur constantly in both directions between time *τ* and *τ*_*p*_. However two-way symmetric migration is not always the case in the real data. It is possible to have a weaker migration from *A* to *B* but a stronger migration from *B* to *A*. Here we examine an extreme case: one-way migration between two species. Without loss of generality, migration is assumed to occur only from *A* to *B* with migration parameter 2*M*. This is to make sure that the total number of migrants remains the same. As a comparison, the overall accuracy is 80.11% using test data with two-way migration. Classification of test data with one-way migration drops to 71.36% (i.e. decreases by 9%). Most of the misclassified samples are simulated with 2*M >* 2. For example, classifier classifies 49% of one-way migration data with 2*M* = 6 as “different species” and only 43% of data samples with 2*M* = 10. Therefore, CLADES works reasonably well for the unsymmetrical migration case when the migration level is moderate, but when migration rate is larger, CLADES performs less well.

We also test how classifier performs with population size bottleneck, a more ancient population splitting time *τ*_*p*_ and various recombination rate. For simplicity, bottleneck is assumed to occur at a specific time before species *A* and *B* split. Let *α* denote the parameter of bottleneck and *N*_*o*_ denote the effective population size before and after bottleneck. During time 1.5*θ ∼* 2*θ* in history, bottleneck occurs and population size is *N* = *αN*_*o*_. As shown in Figure 5 (B), accuracy drops by 10% as the bottleneck parameter decreases from 1.0 (no bottleneck) to a severe level 0.2. In addition, our results show that accuracy first increases and then decreases with the increase of recombination rate. Also the classifier is able to delimit species correctly for a more ancient population splitting time *τ*_*p*_ = *τ/*100. Compared to *τ*_*p*_ = *τ/*500, accuracy slightly drops from 80.11% to 79.48% (see the Supplementary Material A for details).

### 3.2 Classification-based species delimitation

#### 3.2.1 Delimiting multiple species

Classification accuracy is tested under various scenarios. Here we demonstrate our classification-based method CLADES can work for multiple species (see section 2.4 for details) by examining the cases with more than two species. Four species are simulated and each species are assumed to have two populations. Figure 6 presents two population tree topologies that are used for the test of CLADES. In the first case, four species have a symmetric topology, species divergence time are *τ*_0_ = *θ* = 0.01, *τ*_1_ = *θ/*2 = 0.005 and population splitting time is *τ*_*p*_ = *θ/*500 = 2 *×* 10^−5^. In the second case, we assume asymmetric topology. Parameters used are *τ*_0_ = *θ* = 0.01, 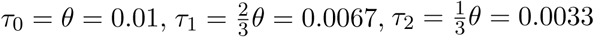 and *τ*_*p*_ = *θ/*500 = 2 *×* 10^−5^ respectively. In both cases, CLADES is able to obtain the maximum likelihood for the correct assignment *{A*_1_, *A*_2_*}, {B*_1_, *B*_2_*}, {C*_1_, *C*_2_*}, {D*_1_, *D*_2_*}*.

**Figure 6:**
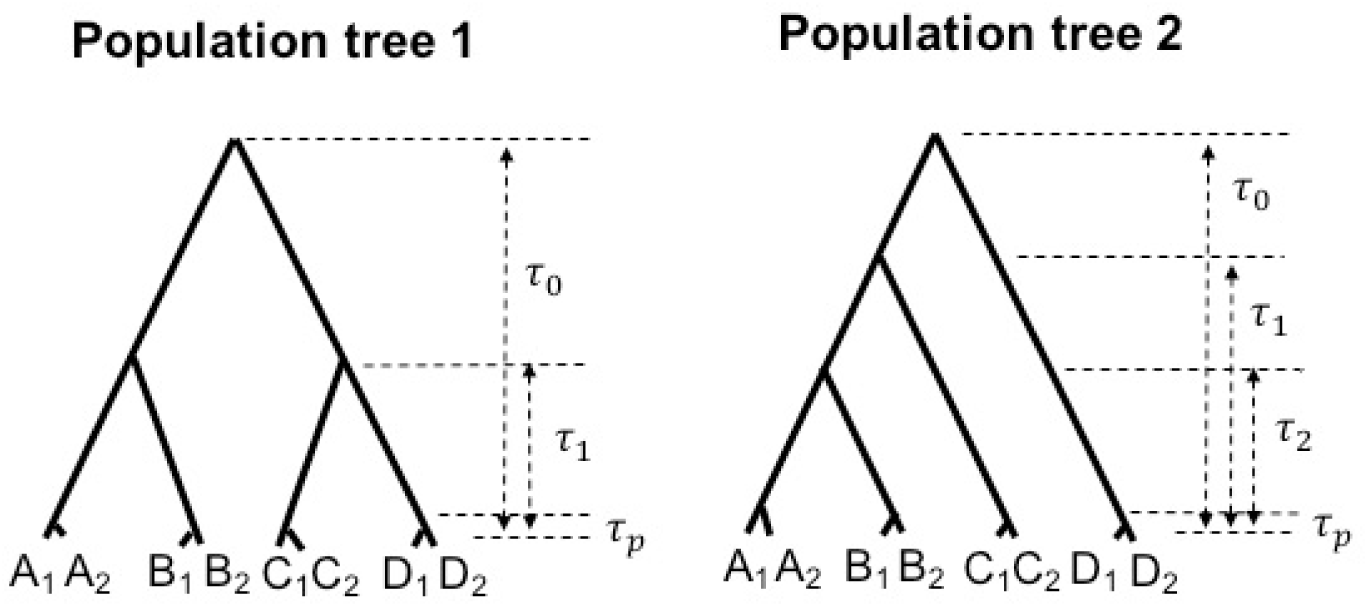
Two population trees for multiple species. There are 4 species and 8 populations in both population trees.

#### 3.2.2 Comparison to BP&P and BFD*

In this section, we compare CLADES with two existing tools BP&P and BFD* that use sequence data and SNP data respectively. To compare with BP&P, we simulate sequence data at 30 loci with length *L* = 2*Kbp* based on two-species model. Population size parameter *θ* and species divergence time *τ* are fixed to be 0.01. For each *M* value from *{*0, 0.01, 0.1, 1, 3, 5*}*, we generate five data replicates. Among these five independent tests, we use the number of tests that BP&P or CLADES suggests the correct species delimitation as the accuracy of the method. BP&P is run for both A10 (guide tree provided) and A11 (guide tree not provided) modes for the same test data (see the Supplementary Material Section B for more details). Figure 7 shows the accuracy of BP&P and CLADES in lines, we can see both methods perform well when *M ≤* 0.1. As *M* gradually increases, more tests of BP&P fail to delimit two species while CLADES only has one test failure when *M* = 5. The box plots shown in Figure 7 (A) and (B) indicate the range of posterior probability for the true species delimitation model estimated by BP&P. The significant decrease of posterior probability and the increase of its variance lead to less accurate delimitation by BP&P. The box plots in Figure 7 (C) show the range of likelihood for the true species delimitation model computed by CLADES. Note that the posterior probability of BP&P and the likelihood of CLADES are not directly comparable. Nonetheless, since these two quantities are the main tools for species delimitation by the two methods, their values should be stable under various settings. As the migration parameter increases, the variance of the likelihood estimated by CLADES appears to be relatively stable in most settings. In contrast, the posterior probability of BP&P appears to have larger variance especially when the migration rate is high. Note that the likelihood shown in Figure 7 (C) is for the optimal species status assignment by CLADES. When *M* increases, this likelihood value decreases. This is to be expected: when *M* increases, CLADES is less certain about the species delimitation.

**Figure 7:**
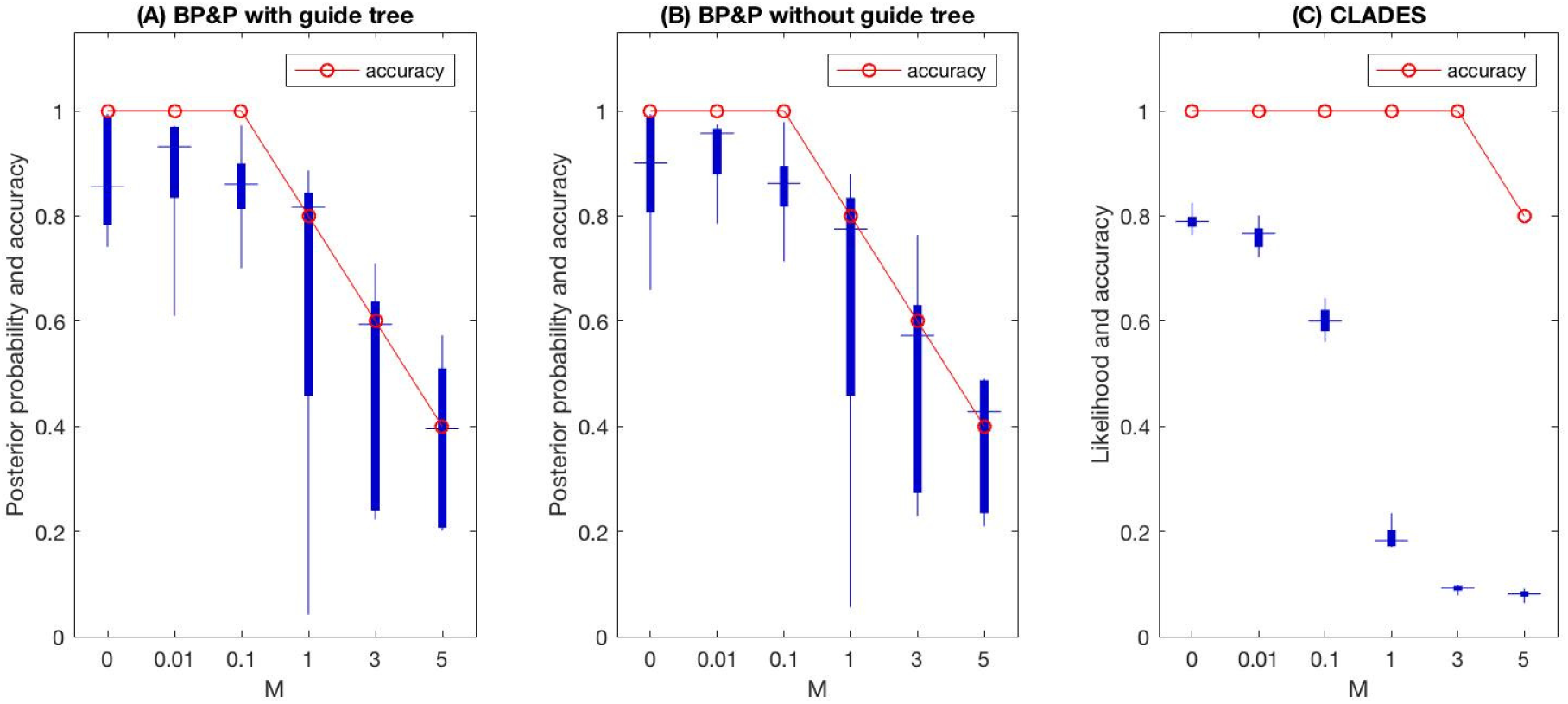
Comparison between BP&P and CLADES. (A) and (B) Box plots for posterior probability of true species delimitation model. (C) Box plots for likelihood estimation of true species delimitation model.

We now compare with BFD*, which uses SNP data. As BFD* is much more time consuming, we only compare two methods on SNP data from one locus simulated with length *L* = 10*Kbp* based on the two-species model for one parameter setting *θ* = 0.01, *τ* = 0.01 and *M* = 1. To run BFD*, four species delimitation models are tested and Bayes Factor is computed against model *{A*_1_*},{A*_2_*},{B*_1_*},{B*_2_*}* (here we use assignment to represent the species delimitation model). More details about priors setting and parameters of MCMC in BFD* are described in Section B of the Supplementary Material. Table 1 contains the estimated marginal likelihood values and Bayes factors for 4 models. The Bayes factors indicate BFD* positively supports model *{A*_1_, *A*_2_*},{B*_1_*},{B*_2_*}* (2 *≤ BF ≤* 10, where BF is Bayes Factor and this indicates strong support). It is possible that the strength of migration prevents BFD* from detecting the species status of *B*_1_ and *B*_2_. Using the same SNP data, CLADES classifies *A*_1_ and *A*_2_ as the ‘same species’ with probability 0.9530 and *B*_1_ and *B*_2_ as the ‘same species’ with probability 0.8066. Thus in this case, CLADES finds the accurate assignment *{A*_1_,*A*_2_*},{B*_1_,*B*_2_*}* with the maximum likelihood.

**Table 1:** Results for 4 species delimitation models from BFD*. Data is simulated at 1 locus from L = 10Kbp based on two-species model. MLE: Marginal likelihood estimate. BF: Bayes factor.

| model | Species | MLE | Rank | BF |
| --- | --- | --- | --- | --- |
| $\{A_1\}, \{A_2\}, \{B_1\}, \{B_2\}$ | 4 | -4648.2912 | 2 | – |
| $\{A_1, A_2\}, \{B_1\}, \{B_2\}$ | 3 | -4643.9139 | 1 | 8.7546 |
| $\{A_1\}, \{A_2\}, \{B_1, B_2\}$ | 3 | -4885.9780 | 4 | -475.37 |
| $\{A_1, A_2\}, \{B_1, B_2\}$ | 2 | -4724.0592 | 3 | -151.54 |

#### 3.2.3 Cryptic sympatric species

We simulate test data with *θ* = *τ* = 0.01 and *M* = 1 based on two-species model, and assume population labels for 40 sequences at each locus are unknown. The clustering method (as described in Section D of the Supplemental Materials) is able to cluster sequences into four groups correctly and then conduct species delimitation with CLADES.

#### 3.2.4 Efficiency

CLADES is very efficient and can scale to large datasets. The running time for CLADES includes summary statistics computation from genetic data, classification of data samples and maximum likelihood estimation. In comparison with BP&P, we use simulated sequence data of 30 loci in length *L* = 2*Kbp* for 4 populations. On average, CLADES takes *∼*2min and BP&P uses *∼*6h to run using one CPU. In comparison with BFD*, 20 individuals and each with 342 SNP sites take BFD* *∼*3h on average for each species delimitation model using 16 threads, while CLADES only takes *∼*1min using one thread.

The running time of CLADES increases linearly with the number of loci in the dataset. If we fix the gene length to be *L* = 2*Kbp* and simulate sequence data with different number of loci, the running time of CLADES increases roughly linearly with the number of loci (see Figure 8). Note that data with 10,000 loci of length *L* = 2*Kbp* only takes 34 min using one CPU. Moreover, CLADES can easily run in parallel with multiple threads as each locus can be first classified independently in CLADES.

**Figure 8:**
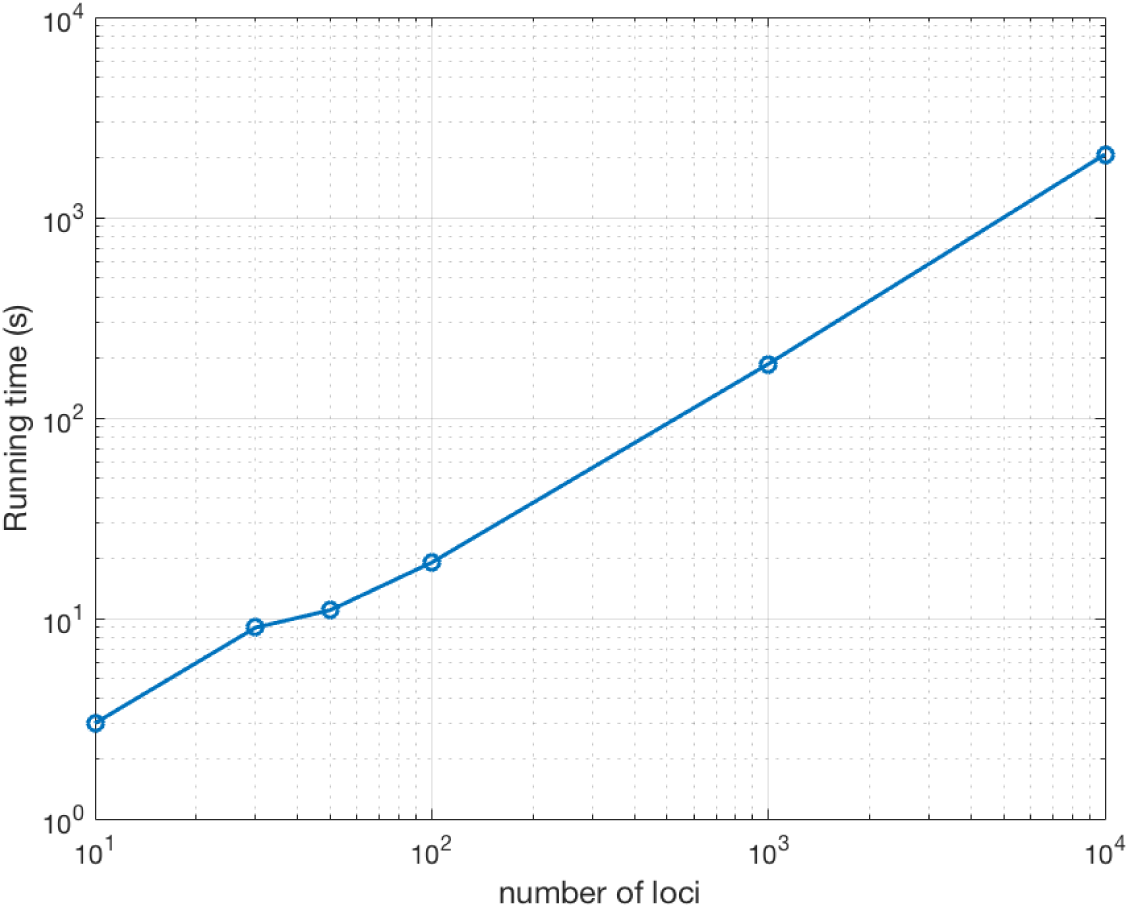
Efficiency on large scale data. Running time of CLADES using different number of loci. To better visualize, x-axis and y-axis use logarithmic scale. Simulation parameters: *θ* = 0.01, *τ* = 0.01, *M* = 0.

### 3.3 Real data analysis

#### 3.3.1 Frog data

We apply CLADES to two related frog datasets obtained from Lu et al. (2014). One dataset consists of three well-recognized frog species *A. lifanensis, A. granulosus* and *A. loloensis* (e.g. Frost et al., 2011). The dataset contains sequence data from two genes, one nuclear diploid gene and one mitochondrial haplotype gene for 19 individuals in total. Although only two loci are available for species delimitation, CLADES strongly supports that *A. lifanensis, A. granulosus* and *A. loloensis* are three different species. Data samples are classified with a probability ranging from 0.8980 to 0.9991 (Table 2). As a comparison, we apply BP&P to the same dataset with three sets of priors (Lu et al., 2014). See Supplementary material section B for details. And the population tree shown in Figure 9 (A) is provided to BP&P. All provided priors suggest three different species, a prior of *θ ∼ G*(1, 10) and *τ ∼ G*(2, 2000) suggests three different species with the highest posterior probability 1.

**Table 2:** Estimation probability based on multiple genes from CLADES for real data. Probability range indicates the range of estimation probability in samples that support a label for pairwise populations.

| model | CLADES |  |
| --- | --- | --- |
| data | different species (+1) | same species (-1) |
| Frog dataset 1 | (0.8980,0.9900) | (0.0099, 0.1019) |
| Anopheles | (0.9948, 0.9976) | (0.0023, 0.0051) |
| Chimpanzee | (0.0221, 0.1803) | (0.8197, 0.9779) |

**Figure 9:**
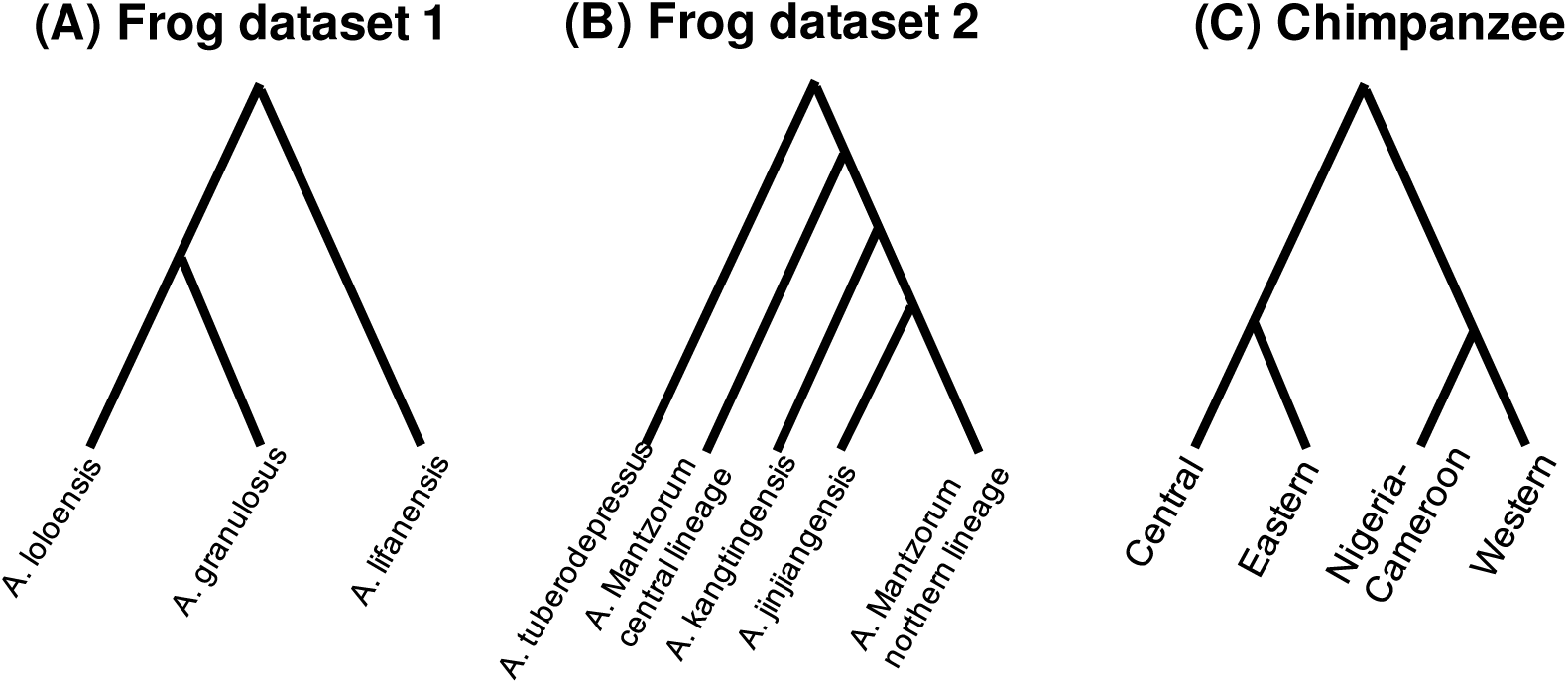
Population tree for real data. (A) Population tree for frog dataset 1. (B) Population tree for frog dataset 2. (C) Population tree for Chimpanzee data.

The other dataset consists of 5 disputable species *A. mantzorum central lineage, A. mantzorum northern lineage, A. kangtingensis, A. jinjiangensis* and *A. tuberodepressus* (Fei et al., 2009), which are related to the three well-recognized frog species described above. How many species there are in this group is under active debate. According to Lu et al. (2014), the GMYC model (Pons et al., 2006) suggests the presence of three different species among *A. kangtingensis, A. jinjiangensis* and *A. tuberodepressus*, but it gives different conclusions for *A. mantzorum central lineage* and *A. mantzorum northern lineage* with different genes. We then apply BP&P to the datasets with the guide tree shown in Figure 9 (B), the phylogeny topology is proposed by Lu et al. (2014). Priors of *θ ∼ G*(1, 10) and *τ ∼ G*(2, 2000) strongly support 5 different species across the populations. However, priors of *θ ∼ G*(2, 2000) and *τ ∼ G*(2, 2000) weakly support a delimitation of two different species *A. tuberodepressus* and others.Note that BP&P results may depend on the choice of priors.

CLADES strongly supports that *A. mantzorum* (including *central lineage* and *northern lineage*), *A. kangtingensis, A. jinjiangensis* and *A. tuberodepressus* are 4 different species. Data from both genes are classified to “different species” with probability mostly greater than 0.8. However, for the disputable species *A. mantzorum central lineage* and *A. mantzorum northern lineage*, the nuclear gene supports they belong to the “different species” with probability 0.9983 but the mitochondrial gene supports they belong the “same species” with a probability 0.6661. This might be due to the significant gene flow (2*Nm* = 0.35) between two populations (Lu et al., 2014). As CLADES is designed to accommodate migration, this level of gene flow between populations are allowed for CLADES. Although CLADES weakly support “different species” (with probability 0.67), the result largely agrees with the GMYC model and the conclusion in Lu et al. (2014).

#### 3.3.2 *Anopheles* data

We apply CLADES to a *Anopheles* dataset including two sibling species from the Itaparica and Florianopolis populations (Rona et al., 2010). The sample consists of 6 genes Clock, Rp49, RpS2, RpS29, cycle and timeless sequences. In the analysis of Rona et al. (2010), *θ* estimated from 6 genes is in the range of 0.008 *∼* 0.043 and the population divergence time is in the range of 1.1 *∼* 3.6*Mya* (estimated to be approximately 2.4*Mya*). In CLADES, all genes support they belong to “different species” with a high probability greater than 0.99 (shown in Table 2). We then delimit species with BP&P for comparison. Since the most frequent estimate of *θ* is around 0.01, we use priors of *θ ∼ G*(1, 100) and *τ ∼ G*(2, 200) in BP&P, and a simple binary tree is provided as the guide tree. Result of BP&P also strongly supports that Itaparica and Florianopolis are two different species.

#### 3.3.3 Chimpanzee from Great Apes

The Great Ape Genome Project (Prado-Martinez et al., 2013) database contains SNP data of four populations of Chimpanzee. Whole genome data are available for 25 individuals from these 4 sub-species, 10 from *Nigeria-Cameroon chimpanzee*, 6 from *Eastern chimpanzee*, 4 from *Central chimpanzee* and 5 from *Western chimpanzee*. The evolutionary history for 4 populations is shown in Figure 9 (C). We use the heuristic method as described in Section 2.3 to compute the summary statistics from unphased genotypes. Chromosome 1 of four populations of Chimpanzee are used for analysis. We use a fixed window length *L* = 10*Kbp* to cut chromosome 1 into pieces. If we regard each piece as a gene, then we collect SNPs within each gene and only use the one for further species delimitation if there are more than 250 SNPs in this gene. After filtering, 213 genes are left for use. CLADES correctly classifies all pairwise populations with a high probability greater than 0.8 (Table 2) and strongly supports that four sub-species are the same species. For further analysis, we select one gene to run BFD* and examine 5 models (see Section B of the Supplementary Material for priors and other settings in BFD*). As shown in Table 3 Bayes factor is computed against species delimitation model of four different species. Clearly, BFD* is strongly against the model of four different species. The model of 1 species (four populations belongs to the same species) present the highest maximum likelihood estimation.

**Table 3:**
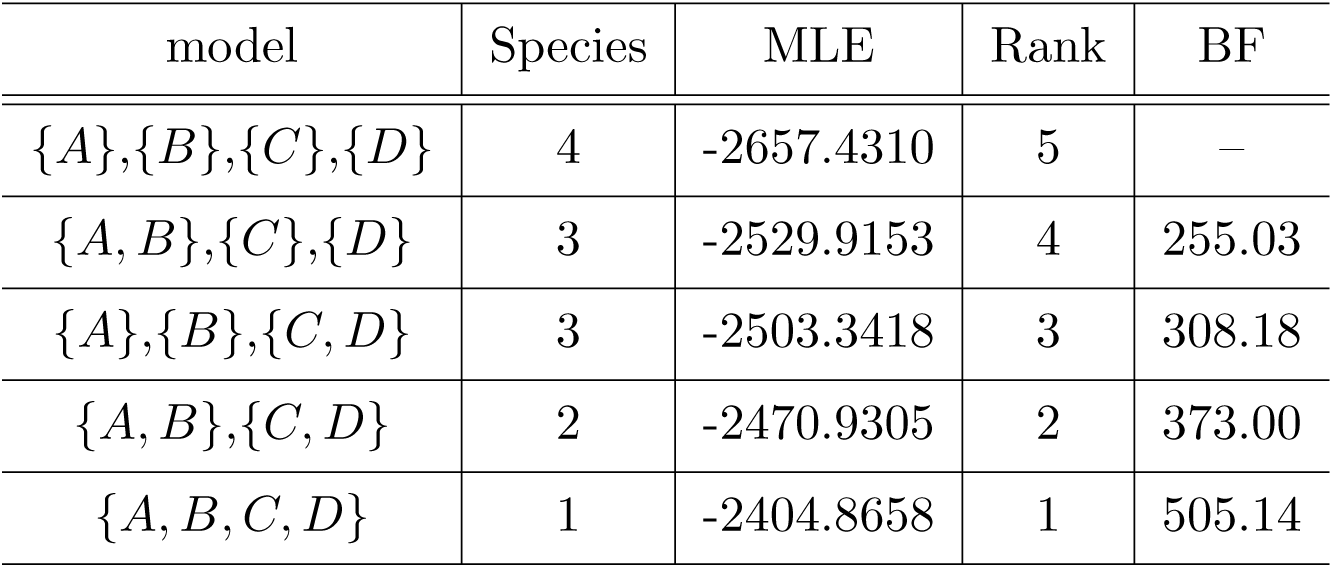
Species delimitation for Chimpanzee populations with BFD*. Here we use letters to represent 4 Chimpanzee sub-species. A: Central Chimpanzee, B: Eastern Chimpanzee, C: Nigeria-Cameroon Chimpanzee, D: Western Chimpanzee. MLE: Marginal likelihood estimate. BF: Bayes factor.

## 4 Discussion

In this paper, we develop a classification-based strategy to delimit species with a broad range of *θ, τ* and *M*. We simulate sequence data based on a two-species model with various parameter settings to train a SVM classifier. As expected, the percentage of accurately classified data samples with different parameters are 80% on average. As we consider a broad range of *θ ∼* (0.0005, 0.02), it is possible to classify two groups of individuals with unknown *θ*, even as small as 0.0005. Moreover, our results indicate that it is desirable to use longer loci to compute summary statistics, especially in the case of more recent divergence (*τ* = *θ/*10). When the migration parameter *M <* 1, data from two species can be classified with accuracy more than 80%. As *M* increases from 1 to 5, CLADES can still classify two groups of data reasonably well with an accuracy higher than 60%. This largely extends the range of migration parameter *M* that Bayesian methods can work with.

Species are the basic unit of biological studies. However, there are many disputable/cryptic species in the nature. Some of species are hard to delimit based on traditional morphology and genetic methods, especially for those of more recent origin. Blurred species boundaries confuse biologists and impede the progress of our knowledge and science seriously. Previous methods of species delimitation need prior knowledge of *θ, τ* and even guide tree. However, users-provided priors could be wrong when prior knowledge about these disputable species is absent. In fact, this situation is likely not rare. Furthermore, in practice, population size parameter (*θ*) and migration rate (*M*) vary in different species. Our simulation indicates that these two parameters can heavily affect the accuracy of species delimitation when using previous methods.

One issue with CLADES is on the selection of summary statistics. The choice of summary statistics depends on whether it can help to divide data samples into two clusters. For the summary statistics we used in this paper, they all show high correlation with cluster label (“same species” or “different species”). And the five summary statistics used in this paper can work reasonably well in the classification of samples. One reason is that these five summary statistics can be computed for both sequence and SNP data and can capture the level of genetic differences between two populations and the level of migration between populations. Moreover, CLADES allows simple customization by adding more summary statistics. One can add more summary statistics as long as it helps in classification. For example, suppose a phenotype is important for some specific species, then one may add the phenotype as another summary statistics and train a customized classifier instead.

*F*_*st*_ is an important summary statistic. *F*_*st*_ and the pairwise difference ratio we defined both have a high correlation with cluster label (“same-species” and “different-species”). According to their definition, they are computed using the same information. However, they are not computed in the same way and thus both are used as features. In other words, they do not have a linear relationship. If we denote pairwise difference ratio as *X*, then *F*_*st*_ would be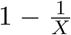. On the one hand, two summary statistics with non-linear relationship won’t affect the stability of classification. On the other hand, using one more summary statistics in SVM is to divide the space into two by searching in one more dimension. For example, in the extreme case when the pairwise difference ratio is close to 1, a *F*_*st*_ *<* 0 would suggest it is more likely to be “same-species”. And in training we do observe the increase of more than 3% in accuracy when adding *F*_*st*_ as another feature. Since the boundary of summary statistics to divide space is not strictly defined, it would be better to use multiple planes (i.e. multiple features) in classification.

Existing species delimitation approaches are either too slow to handle genome-scale data or do not account for gene flow. For example, BP&P are slow for large data and also do not consider accommodate gene flow. BFD* can use genomic SNP data but cannot account for gene flow. PHRAPL can account for gene flow but does not easily scale up to genomic data.

CLADES, a machine-learning approach, is guide tree free and prior free method that can work with both sequence data and SNP data. CLADES also works well when the *M* value is moderate. CLADES is flexible to be extended and designed for specific cases. For example, it is possible to add more useful information (such as phenotype) to train a customized classifier with prior knowledge of closely related species. Furthermore, CLADES is efficient and can be applied to genomic-level data. In summary, CLADES can help biologist to perform species delimitation more accurately and efficiently.

## Acknowledgements

We thank Laura Kubatko for useful discussions. JP, CC, XL and YW are partly supported by U.S. National Science Foundation grants IIS-1526415 and CCF-1718093. BL is partly supported by National Natural Science Foundation of China (Grant No. 31401961) and West Light Foundation of the Chinese Academy of Sciences (Y5C3021100).

## Data Accessibility

Frog dataset is accessible in GenBank Accession Nos. KJ008104KJ008461. Anopheles dataset is accessible in Rona et al. (2010). Chimpanzee data can be downloaded from Great Ape Genome Project website: http://biologiaevolutiva.org/greatape/data.html. CLADES models and code can be downloaded: https://github.com/pjweggy/CLADES.

