## Supplementary Materials for "CLADES: A Classification-based Machine Learning Method for Species Delimitation from Population Genetic Data"

#### A Robustness of classification

In training and testing, we simulate two populations in the same species splitting at time $\tau_p = \tau/500$. However, $\tau_p$ cannot be determined by a specific range. Here we examine a more ancient population splitting time $\tau_p = \tau/100$ to see if the trained classifier is capable to detect this more difficult "same species" case. As a comparison, the overall classification accuracy is 80.11% for test data with $\tau_p = \tau/500$. For each candidate $(\theta, \tau, M)$, we simulate test data using $\tau_p = \tau/100$. The overall accuracy of classifier is 79.48%. The results indicate the trained classifier is able to classify simulated data with a reasonably large population splitting time.

For the simplicity of simulation, We do not consider recombination in the process of generating training data. Here we evaluate how recombination affects the accuracy of classification. Here we fix parameters $\theta = \tau = 0.01$ and $M = 1$, and then vary recombination parameter $r = 4N\rho$ from 0 to 0.02 with interval 0.005. For the recombination parameter value in {0,0.005,0.01,0.015,0.2}, the classification accuracies are 73.33%, 76.11%, 83.89%, 91.11% and 86.67%, and F-scores are 0.76, 0.78, 0.86, 0.92 and 0.89. The accuracy first increases and then decreases which indicates recombination can help to distinguish two populations at certain level.

In addition, as *MCcoal* is limited in simulation of recombination and bottleneck in history, we use another simulator *ms* by Hudson (2002) to simulate test data for evaluation of impact of recombination and bottleneck.

#### B BP&P and BFD* settings

In Section 3.2.2, we compare CLADES with two existing tools BP&P and BFD*. When using BP&P, two species delimitation modes are applied: one is to provide guide tree topology (A10 mode) and the other does not require guide tree (A11 mode). For each test, we use Gamma distributions with the mean equal to the true value of $\theta$ and $\tau$ as priors. Note that often we don't have such ideal settings for real data analysis. For example, if test data is simulated with $\theta = \tau = 0.01$, then we use priors $\theta \sim G(1, 100)$ and $\tau \sim G(1, 100)$ in running BP&P. In addition, we set burn in with 4000 and 200000 samples are generated. In analysis of frog datasets with BP&P, three sets of priors examined are: (1) both $\theta$ and $\tau$ follow $G(1, 10)$ (2) $\theta \sim G(1, 10)$ and $\tau \sim G(2, 2000)$ (3) both $\theta$ and $\tau$ follow $G(2, 2000)$ (Lu et al., 2014).

When running BFD*, the default usage of mutation rate calculation is used to decide the mutation rate of the data. Similar with BP&P, we use the Gamma distribution with mean equal to the true value and also follow suggestion by the table in the tutorial of BFD* to decide Lambda value. For stability of data sampling, we use at least 48 steps, chainLength=100,000 and preBurnin=10,000 in MCMC setting. In analysis of Chimpanzee dataset with BFD*, we use the same MCMC setting and a prior $\theta \sim G(2, 2000)$ according to the conclusion of Prado-Martinez et al. (2013). To obtain the maximum likelihood estimation for species delimitation model of a single species, we add one human individual as out of group genotype because BFD* cannot directly compute maximum likelihood for all individuals falling into the same species.

### C Heuristic method for computing longest shared tract for genotype data

When genotype data is not phased to haplotypes, we propose a heuristic method for computing the longest shared tract with genotype data. Note that each diploid individual has a genotype which is composed of two haplotypes. We want to estimate the length of the longest shared tract between haplotypes of two individuals from two unphased genotypes. As a concrete example, we consider a pair of genotypes as shown below. Here we use 0/1/2 to denote genotype data. 0 and 2 are the homozygotes and 1 is the heterozygote. For example, the first site has (0, 0), and the second site has (2, 2) but the third site presents (2, 0). The first two genotypes at the same SNP sites are (0, 0) and (2, 2), which indicate that such sites must be 'shared' (i.e. same). However, genotypes of (0, 2) or (2, 0) indicate that such sites must not be 'shared' (i.e. different). In case the genotype is 1, it is possible that one haplotype is shared between two individuals. Thus we first consider all (0, 0), (2, 2), (1, $\cdot$) and ($\cdot$, 1) as 'shared' to compute the upper bound of longest shared tract (where $\cdot$ can be 0/1/2). Then we compute the lower bound of longest shared tract by regarding the first (1, $\cdot$) and ($\cdot$, 1) encountered as 'shared' but the second (1, $\cdot$) and ($\cdot$, 1) encountered as 'not shared'.

02201002102
02010002012

Note that the position for each SNP site is known. As we move along the SNP genotypes, we can compute the length of shared tract in base pair resolution. Then we take the average over the upper bound and the lower bound of longest shared tract as the longest shared tract for two unphased genotypes.

### D Analysis for cryptic sympatric species

So far we assume we are given population labels of all samples. Sometimes this is not the case in the field. Sometimes a group of organisms are collected where these samples are from different populations or species. When dealing with data that are arbitrarily grouped together by sample locality, genetic populations are not well-defined. One more step is required before using CLADES for species delimitation. Suppose there are $n$ sequences collected by sample locality and $k$ potential populations across the data. We assume the number of populations $k$ is known. We can use K-means clustering (Hartigan and Wong, 1979) to cluster $n$ sequences with $k$ clusters by using the Hamming distance ($n \geq k$). We then treat the $k$ clusters as the populations, which are used for further species delimitation analysis. We simulate test data of one locus with length $L = 10Kbp$ and using parameters $\theta = \tau = 0.01$ and $M = 1$ based on the two-species model. Suppose these sequences are collected by sample locality, and so these sequences are in a single group. We use the K-means method to cluster 40 sequences to 4 clusters. Result shows that clustering method is able to detect 4 populations successfully. Therefore it is feasible in some cases to delimit cryptic sympatric species from mixed-up samples.
